# LoopBin – a VaDE-based neural network for chromatin loop classification

**DOI:** 10.64898/2026.01.13.699359

**Authors:** Yajie Zhu, Argyris Papantonis, Mariano Barbieri

**Affiliations:** Institute of Pathology, University Medical Center Göttingen, Germany

**Author notes:** Correspondence to: A.P.;, M.B.

**Keywords:** chromatin folding, Hi-C, Micro-C, histone marks, enhancer-promoter looping, CTCF, genome topology, machine learning

## Abstract

Classifying chromatin loops from 3D genomics data according to their epigenetic and structural attributes is important for inferring their functional roles. Currently, such classification typically relies on the manual intersection of epigenomic signal peaks and loop anchor locations. To automate this, remove peak-calling biases and include information inherent to 3D genomics signal structure, we developed LoopBin, a framework based on a variational deep embedding (VaDE) neural network. We applied LoopBin to kilobase-resolution Micro-C data and segmented tens of thousands of loops into clusters with distinct features using minimal histone modification and transcription factor-binding data. These features were indicative of apparently distinct biological function by each subgroup of loops. Therefore, LoopBin can provide insights into the dynamic shifts in loop classification that can occur upon perturbation of cell homeostasis or signaling.

## INTRODUCTION

Chromatin loops, defined as point-to-point 3D interactions between genomic loci, constitute an important layer of genome organization inside the nucleus. For example, loops can bring distant *cis*-regulatory elements—such as enhancers or silencers—into spatial proximity to their target promoters to exert gene activation or repression (1). As such, many different types of loops are likely to form along metazoan chromosomes, and loops may disappear, form *de novo* or rewire as the cellular context changes (2–6). Currently, loop type annotation predominantly relies on matching loop anchors with signal peaks called from histone marks (e.g., H3K4me3 typically marking gene promoters) or chromatin-binding factor ChIP-seq profiles (e.g., CTCF typically found at both anchor points of ‘structural’ loops; 7). However, there are obvious caveats to this approach: the overlap with ChIP-seq peaks is a binary readout affected by empirical background and enrichment thresholding; loop anchors are often marked by overlapping signals of variable intensity; differences in the structure of the loop interaction signal itself are not considered.

Classical clustering approaches cannot be readily applied for the annotation of chromatin loops due to issues with the high dimensionality and noise levels in 3D genomics (e.g., Hi-C or Micro-C; 8) and ChIP-seq (or similar) data. To date, ‘deep’ learning models have been used in 3D genome organization studies for predicting loop formation (9–11) or for imputing low resolution contact maps (12–16) by effectively reducing dimensionality and capturing complex non-linear relations in the data.

To address this gap, we opted for a variational autoencoder architecture, VaDE, a deep learning method for dimensionality reduction that simultaneously identifies data clusters, outperforms classical clustering methods, and is considered a standard approach in the field of deep clustering (17–20). The typical autoencoder (AE) architecture aims at representing observed data at lower dimensionality (21–23). The initial encoder part transforms a high-dimensionality input into a low-dimensionality embedding termed ‘latent space’, while the decoder part reconstructs the input from this latent embedding. By minimizing the difference between the reconstructed output and the input during training, the model learns the most significant features that can be used for clustering (reconstruction). In addition, by minimizing on a mixture of Gaussian priors (GMM) in the latent space, the VaDE architecture interprets the lower-dimensional representation in terms of different clusters (regularization process minimizing the Kullback-Leibler (KL) divergence; 23). By jointly optimizing reconstruction and regularization loss functions, a continuous cluster-friendly latent space is learned (17,23).

We exploited VaDE architecture to build the LoopBin model, a neural network for the automatic classification of chromatin loops and their corresponding epigenomic profiles into more intricate, structure-aware subgroups. We trained LoopBin using loop-rich Micro-C from human cells (24) and matching CUT&Tag epigenomic data (25). LoopBin classified Micro-C loops into clusters with distinct biological attributes that manual classification would not reveal. These attributes are robust and dependent on the provided epigenomic profiles. Simulations of chromosome folding further confirmed that LoopBin identifies loop classes associated to mechanistically different 3D chromatin interactions. This exemplifies the usefulness of our framework in the continuous effort to understand the functional purpose and dynamic nature of loop formation in different contexts.

## RESULTS

### Overview of the LoopBin framework

In LoopBin, we adapted the VaDE architecture to integrate and analyze matched Micro-C and CUT&Tag data (**Fig 1A,B**): an encoder-decoder architecture transforms the combined input data in a latent space representation; a mixture of *k* Gaussian priors in the latent space—with mean μ*_k_*, variance θ*_k_* and prior probability π*_k_*—identifies the clusters (17). Next, a pre-training phase on a simpler AE architecture is run to learn the starting weights of the encoder and decoder for the training phase and prevent trapping in local minima, which commonly affect variational autoencoders (17,19,20). In addition, we modified the pre-training phase to learn μ*_k_* and θ*_k_*, and use them as hyperparameters in the training phase. This modification preserves training stability by avoiding GMM collapse to a unit Gaussian and improves cluster separation in the latent embedding (**Fig S1A**). The number of clusters in the mixture is selected by the user and fixed. Remarkably though, the effective number of clusters converges during the learning process by the spontaneous emptying of loops in redundant clusters (π*_k_* decreases to zero in these clusters). This suggests that LoopBin captures inherent features in the data independent of the initially selected cluster number (Fig S1B).

**Figure 1.**
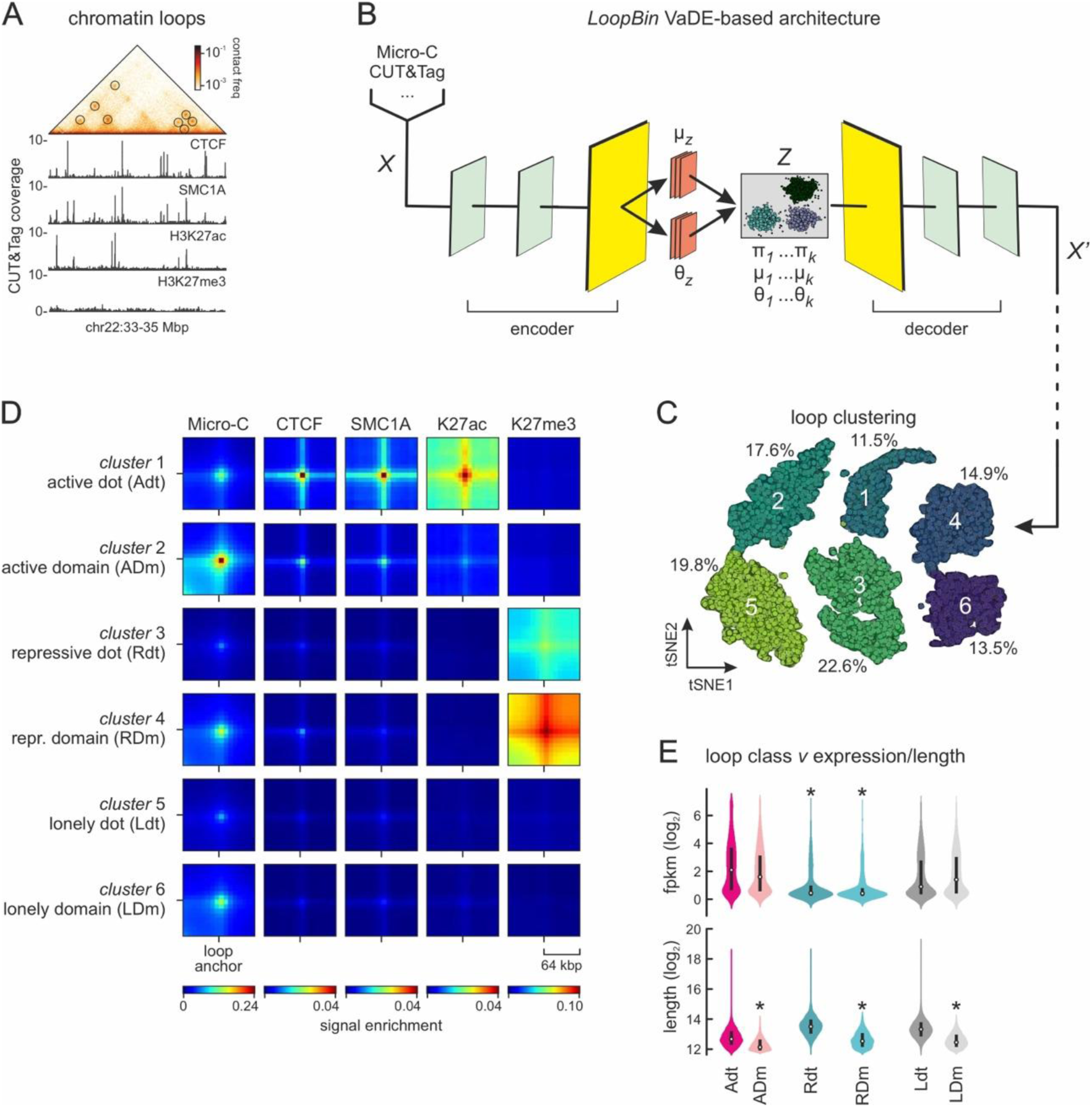
LoopBin classification of Micro-C chromatin loops and epigenetic marks. **A**, Exemplary views of the data used as input for LoopBin: Micro-C contact maps and CUT&Tag occupancy profiles for CTCF, SMC1A (cohesin), H3K27ac and H3k27me3. **B**, LoopBin architecture: an encoder transforms input *X* into a latent space distribution represented by the mean μ*z* and the variance θ*z*. A decoder reconstructs *X*’ from *z*, which is sampled from the latent distribution. *Z* is limited to the Gaussian mixture field represented by the prior probability π*k*, mean μ*k*, and variance θ*k*. **C,** t-SNE plot showing the latent space of loops separated into six clusters. **D**, Average pileup of Micro-C and multiplied CUT&Tag signal between the anchors of loops in each cluster. **E**, Violin plots showing the expression level of genes assigned to loops (top) and the length of loops in each LoopBin cluster.

As input *Χ* for LoopBin, we used concatenated data. First, we took 16×16 windows centered around each of the 44,828 chromatin loops identified in our Micro-C data (24), where each bin is 8 kbp in breadth. These loops include all types of manually annotated interactions, e.g., those involving enhancers and promoters (E-P), enhancers only (E-E) or CTCF-bound anchors. Each Micro-C loop signal was linearized to give a 16×16=256 vector. Second, we transformed genome-wide CUT&Tag profiles from two histone modifications—H3K27ac and H3K27me3, from the loop anchoring CTCF protein, and from the loop-extruding cohesin complex component SMC1A (from ref. 25) into 16-bin vectors. In doing this, we avoided conventional peak calling, but rather binned CUT&Tag signal into 8-kbp non-overlapping bins. This reduces biases due to arbitrary data processing thresholds, and feeds a richer, more continuous signal into LoopBin. Finally, 16-bin CUT&Tag vectors overlapping loop anchors were concatenated with the corresponding 256-bin Micro-C vectors to generate the final input for LoopBin.

### LoopBin annotates loop classes with distinct features

For training and classification predictions, we inputted the concatenated loop and epigenomic vectors into LoopBin. Vectors were deconstructed and the loss function in our VaDE architecture (described in ref. 17) decreased smoothly during training (**Fig S1C**). This ensures that the latent embedding *z* after data reconstruction preserved the most salient features in *X*, while the KL loss regularized *z* to conform to a mixture of Gaussian manifolds. In other words, loops with similar 3D interaction and epigenomic patterns will be assigned to the same cluster in the latent space.

In this case, LoopBin predicted as the most likely outcome the presence of six major loop classes. Each class appeared well-separated from the others (**Fig 1B**), each containing 10-25% of the total number of loops (**Fig 1C**). Thus, all loops are accounted for and annotated without dispersal into small classes populated by few loops with outlier characteristics.

Next, we traced the LoopBin classification back to the structural and signal enrichment features of the underlying Micro-C and CUT&Tag data for each of the six classes. A first observation was that there were two different loop signal structures that LoopBin identified: a ‘dot’ structure indicating a focal interaction between the two anchors, and a ‘domain’ structure where interaction signal arises more diffusely between the anchor boundaries (**Fig 1D**). A second observation was that three major epigenomic profiles were assigned to the six groups: two groups enriched for the ‘active’ H3K27ac and depleted of the ‘inactive’ H3K27me3 histone mark, two groups showing the converse pattern, and two showing lower level CTCF/cohesin demarcation only. As a result, we named these loop classes ‘active dot’ (Adt) and ‘active domain’ (ADM), ‘repressive dot’ (Rdt) and ‘repressive domain’ (RDM), and ‘lonely dot’ (Ldt) and ‘lonely domain’ (LDM), respectively (**Fig 1D**). This LoopBin classification largely agreed with a manual annotation of loop anchors (**Fig S2A,B**), which cannot discriminate between ‘dot’ and ‘domain’ loops though as it does not account for differences in Micro-C signal structure. Interestingly, the discrimination between ‘dot’ and ‘domain’ loops is linked to loop size; longer-range loops are consistently overrepresented in the ‘dot’ class, while shorter-range ones present predominantly as ‘domains’ (**Fig 1E**).

Furthermore, ‘active’ loop classes show H3K27ac enrichment on at least one anchor, indicative of an active regulatory *cis*-element—enhancer or promoter—involved in a presumably regulatory 3D interaction. Here, the strong cohesin signal enrichment is in line with the notion that longer-range ‘dot’ loops rely more on extrusion for their formation and maintenance (1). In contrast, ‘repressive’ loop classes are depleted of cohesin signal in line with the documented antagonism between extrusion and H3K27me3-marked chromatin domains (26). Interestingly, ‘repressive’ loops seem to have higher H3K27me3 on the outside of their anchors (**Fig 1D**). When combined with RNA-seq data (i.e., by assigning gene promoters ±5 kbp to loop anchors), there were significantly more promoters associated with ‘active’ than with any other loop class anchors (65% for ‘active dots’, 50% for ‘active domains’ compared to <40% in the other classes; **Fig S2C**). Accordingly, ‘active dot’-associated promoters showed highest expression, followed by ‘active domain’-associated ones, while genes linked to ‘repressive’ classes were the least expressed (**Fig 1E**). Thus, LoopBin classification reflects biologically meaningful types of 3D chromatin interactions.

### LoopBin classifications are both robust and plastic

For testing classification robustness, we prefixed the number of clusters to 10 and performed five independent LoopBin runs. The framework consistently returned 5-7 loop classes with large overlap between runs. For instance, in the run where seven loop classes were returned, the framework divided the larger ‘active dot’ class (**Fig 1D**) into two smaller classes with higher or lower H3K27ac signal (**Fig S3A**). Overall, the loops included in matched classes resulting from the different runs remained essentially the same, with rare instances of a shift from an ‘active’ to a ‘repressive’ class or *vice versa* (**Fig S3B**). Shifts between homotypic ‘dot’ and ‘domain’ classes were more frequent, likely due to the continuous spectrum of Micro-C signal patterns (**Fig S3B**). We also calculated pairwise normalized mutual information (27) to quantify the similarity between different runs and got medium-to-high scores (**Fig S3C**). Thus, LoopBin largely preserves loop assignments over runs, but is also responsive to the inevitable continuity of the data signal it is trained on.

Next, we tested whether the number of inputted epigenomic profiles affects classification. We added H3K4me1 CUT&Tag data, a mark of potentially active enhancers, from DLD-1 cells (28) to the LoopBin input. Addition of this mark did not change LoopBin reconstruction power (**Fig S4A**), but did increase the number of returned loop classes from six to seven (**Figs 1** and **S4B**). Visualization of signal enrichment from Micro-C and each of the five epigenomic datasets for each loop class showed that the six classes predicted before remained, but a new one now emerged: 898 loops, with features likening this of the LDt class, now showed medium H3K4me1 enrichment at their anchors and were separately classified as ‘enhancer domain’ (eDM) loops (**Fig S4B**). The lack of H3K27ac enrichment at eDM loops suggests that they may represent 3D contacts involving ‘latent’ enhancers (29). As would be expected, H3K4me1 also marked the anchors of ‘active dot’ and ‘active domain’ loops, but without changing these classes (**Fig S4B**).

Conversely, when we only used Micro-C and H3K4me1 data as input for LoopBin, just three loop classes were predicted (despite the maximum cluster number being set to ten; **Fig S4A**), which reflected a mere correlation of Micro-C interaction strength with H3K4me1 levels (**Fig S4C**). This shows that LoopBin is best used when mulitple epigenomic datasets are available alongside 3D interaction data, and the increase in the number of input datasets will result in an increasingly nuanced loop classification. At the same time, LoopBin does not overfit and consistently predicts a limited number of classes, even when few datasets are fed.

### LoopBin classifications reveal different 3D chromatin interaction modes

Formation and dissolution of chromatin loops occurs via multiple, often competing mechanisms (1). Therefore, we wanted to understand whether the different types of loops predicted via LoopBin reflect different modes of 3D interactions. Especially in the case of homotypic ‘dot’ and ‘domain’ loops, this may help clarify what their differences are from a standpoint of chromatin folding mechanics.

To this end, we turned to our established molecular dynamics simulation framework (24,30). In these simulations, proteins known to mediate chromatin looping, like CTCF/cohesin or RNAPII, are modeled as beads binding chromatin at experimentally defined rates (see **Methods**), but giving rise to loops via different mechanisms. Cohesin will load onto chromatin and actively extrude loops until it encounters a CTCF-bound chromatin site oriented against the direction of extrusion (7), while RNAPII will directly bridge two or more promoters and/or enhancers via microphase separation (31). These mechanisms co-exist in our simulations and act synergistically or competitively based on the genomic context. The basic assumption here is that all loop anchors in a given class will share the same mode of interaction due to the mediation of similar protein species, thereby producing loops of similar contact and epigenomic signal architecture.

We modeled the whole of human chr21 at 10-kbp resolution with our basic model not allowing for interactions between loop anchors from different classes. We populated the full chromosome length with anchors from the six classes (**Fig 1D**) in addition to CTCF binding sites from CUT&Tag data, where cohesin-mediated extrusion would stall (**Fig 2A**). This produced contact maps that fit well to actual Micro-C data from that chromosome and required both loop extrusion (LE) and microphase separation (PS; **Fig 2A,B**). With this in hand, we asked whether the ‘dot’ versus ‘domain’ structures that LoopBin predicts are merely a result of anchor separation on the linear chromosome. We merged the ‘active dot’ and ‘active domain’ classes in a modified model, so that they arise via the same protein species. Comparing the results of this ‘active-merged’ model to those from our basic one, we saw that new cross-class loops emerged that were not found in actual Micro-C maps (**Fig 2C**). This suggests that the ‘dot’ versus ‘domain’ distinction revealed by LoopBin is due to more than a difference in linear anchor separation.

**Figure 2.**
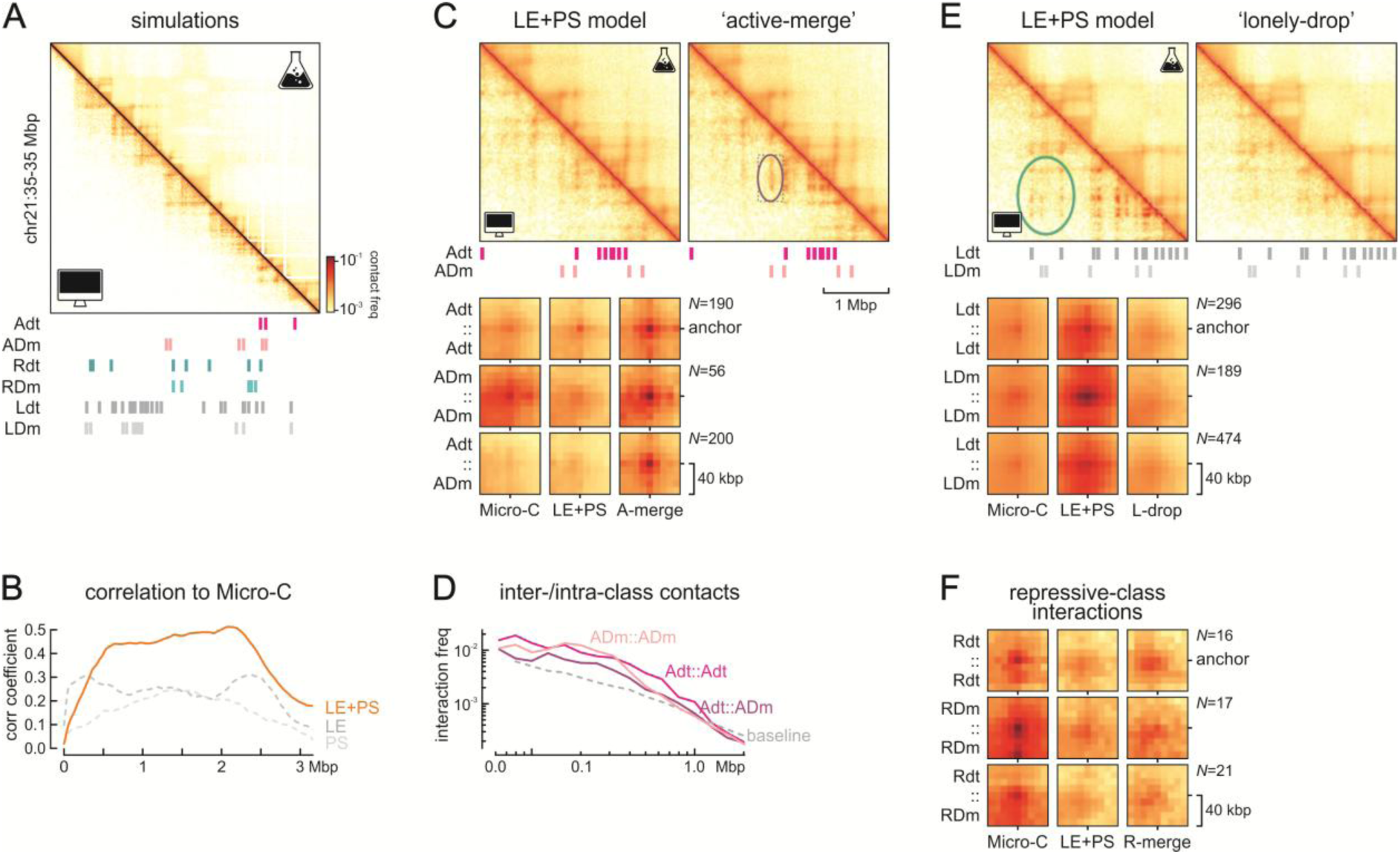
LoopBin informs on chromatin 3D interaction modes. **A**, Comparison of contacts from Micro-C and from Molecular Dynamics simulations of chr21 folding in a 10-Mbp region at 10-kbp resolution. **B**, Correlation by genomic distance between Micro-C maps and simulations (PS—phase separation, LE— loop extrusion, LE+PS—loop extrusion plus phase separation plus loop extrusion). **C**, Top: The ‘active-merge’ model produces inter-active (Adt-ADm interactions) class contacts not observed in Micro-C data. Bottom: APA plots from Micro-C and simulations using the basic or ‘active-merge’ parameters. **D**, Inter-active class (Adt-Adm) interactions are weaker than intra-active (Adt-Adt, ADm-ADm) class ones, but stronger than expected by reference interaction (dashed line). **E**, As in panel C, but for Ldt-LDm class interactions. **F**, APA plots from Micro-C and simulations using the basic or ‘repressive-merge’ parameters.

At the same time though, ‘active domain’ loops appeared stronger than ‘active dot’ ones in this ‘active-merged’ model, which compared better with actual Micro-C profiles than the basic model did (**Fig 2C**). We therefore plotted Micro-C interactions over distance for the different loop classes to discover mild inter-active class interactions (**Fig 2D**). These were not as strong as intra-active ones, but significantly stronger than expected by chance (i.e., by interactions between pairs of genomic segments >80 kbp away from anchors in each respective class; **Fig 2D**). Thus, for loops with active histone mark demarcation inter-class contacts can form, albeit at lower frequency.

Next, we looked into ‘lonely’ loop classes. These loop anchors displayed weak enrichment for cohesin/CTCF (**Fig 1D**), and their interaction strength in Micro-C data was far less that in our basic simulation model (**Fig 2E**). To remedy this, we removed any specific interaction between the protein species binding ‘lonely’ anchors, and only allowed extruding cohesin transversing the chromosome to stall at these positions when bound. This modified ‘lonely-dropped’ model reduced interaction strength within this loop class and now closely recapitulated Micro-C profiles (**Fig 2E**), which hints to a different, indirect mechanism underlying this subset of loops.

Finally, we looked into the ‘repressive’ loop classes (**Fig 1D**) that showed strong inter- and intra-class interactions in actual Micro-C data, which were not well represented in our basic simulations (**Fig 2F**). Upon merging of the two repressive loop classes, contact patterns reflected intra and inter-class interactions more evenly and matched Micro-C profiles better (**Fig 2F**). This suggests that repressive domains differ to active ones in how they give rise to loop structures, especially as regards interactions between loops—and this aligns with current understanding of H3K27me3-marked higher-order domains (26,32–34). Together, the combination of simulations and LoopBin classification can unveil intricacies in the features that underlie loop formation and maintenance, inform the adjustment of modeling parameters and ultimately discriminate between principles of architectural and regulatory domain formation along chromosomes.

## Discussion

The advent of machine learning and neural networks for the processing and analysis of genomics datasets is quickly transforming genome biology (35–37). We exploited the advantages offered by VaDE architecture to produce a framework for the integration of 1D epigenomics data with loops structures from 3D genomics experiments in order to execute automated annotation of loop classes. Our framework, LoopBin, robustly predicts the same type of loop categories in different iterations, with any variability reflecting the continuity of the input data signals. This is, however, important as it alleviates biases from conventional peak calling that essentially binarizes the data. Moreover, LoopBin is scalable (can integrate any type and number of 1D datasets) and applicable to loops derived from any technology mapping genome-wide 3D chromatin interactions (e.g., Hi-C, HiChIP, (Promoter-)Capture-HiC, Micro-C).

Here, we have used the output of LoopBin to inform simulations of 3D chromatin folding. For instance, we deduced that loops marked by H3K27ac may not form by compatible mechanisms so as to produce strong interactions between ‘dot’ and ‘domain’ classes. In contrast, loops marked by the H3K27me3 repressive modification are compatible and interactions between ‘dot’ and ‘domain’ classes can form. These insights do not only allow us to refine simulation frameworks, but also uncover potential mechanisms underlying loop-like chromatin structures that merit further experimental investigation.

We envisage LoopBin being useful for both automated loop classification in different contexts, but also as a framework that can be applied to uncover relationships between loop classes and the mechanisms responsible for their emergence. To cite one example, combining LoopBin with data from highly rearranged cancer genomes (4) might potentiate discovery of disease-related classes of 3D chromatin interactions and help understand the roles and selection in malignancy.

## Methods

### The LoopBin framework

Our model consists of an encoder, a decoder, and a probability model. Their architecture is adapted from VaDE (17,18) with the encoder having a 384-500-500-2000-10 input dimension to latent size structure, and the decoder a 10-2000-500-500-384. The LoopBin probability model is based on Gaussian mixture models, with its number of components, cluster center μ*_k_*, and variance θ*_k_* being set as hyperparameters by the user, while the prior probability π*_k_* is set to 1 divided by the number of components. The code for LoopBin is available at: https://github.com/YajieZhu018/LoopBin.

### Data preprocessing and LoopBin (pre)training

Micro-C and CUT&Tag data of DLD-1 cells (25; GEO accession: GSE178593) were used for training and predictions. Genomic loops were identified at 5kb and 10kb resolution using a combination of HiCCUPS (7), *cooltools* (38), and *mustache* (39) and merged with *pgltools* (40). H3K4me1 CUT&Tag data from DLD-1 cells were publicly available (from ref. 28; GEO accession: GSE193648).

To generate the input for the model, a 16×16 submatrix was extracted centered around each loop from 8-kbp resolution Micro-C matrices, with submatrices including the matrix diagonal excluded in order to avoid the influence of strong interaction therein and then dominate what the framework learns during pre-training. Submatrices were next transformed via logarithmic and ‘min-max’ normalization into a 0-1 range across samples. The CUT&Tag datasets were converted from *bigwig* to *bedgraph* format of 8-kbp resolution and split by chromosome for parallelization. A 16-bin region centered around each anchor was extracted for a given loop, transformed via logarithmic and ‘min-max’ normalization in a 0-1 range. Then, each 16×16 Micro-C submatrix is linearized into a 256-D vector and concatenated with all corresponding CUT&Tag vectors into a final 384-D vector.

For pre-training, the autoencoder has the same architecture as the one in LoopBin, and control and degron data were combined as the input. We pretrained via an Adam optimizer with a learning rate of 0.02 and batch size of 256 for 200 epochs, and applied GMM of seven components to the latent space to get intial μ*_k_* and θ*_k_*. Finally, the encoder and decoder were initialized using the weights from the pretrained autoencoder, and the same type of data used for pre-training were inputted with a batch size of 256 for 2000 epochs with the loss function and the learning rate scheduling defined as described previously (17).

### Molecular Dynamics simulations

The entire chromosome 21 was modelled based on a previously described modelling framework (25,30) using the multipurpose EspressoMD package (41). Briefly, the chromatin fiber is modelled as a self-avoiding polymer chain consisting of equisized beads representing 10 kbp of chromatin. Beads were classified as binding CTCF (based on CTCF CUT&Tag profiles from ref. 25, including motif orientation), as binding category-specific factors at loop anchor sites (belonging to the different categories identified by LoopBin), or as neutral (with none of the above-mentioned signatures). Polymers were used in molecular dynamics simulations in a 3D space following Langevin equations to model thermal motion of chromatin and its binding factors in an implicit solvent (the nucleoplasm), with the following postulations: (i) the category-specific factors have interaction affinity (with same intensity for all categories), hence can bridge beads of the same category by spontaneous spatial co-localization forming loops of the given category; (ii) extrusion of loops by cohesin stalls when encountering a CTCF-bound site with a motif oriented towards the direction of extrusion or when meeting a loop anchor bound to a category-specific factor. By dynamically forming and dissolving protein-chromatin bonds, this framework simulates chromatin loop extrusion by the cohesin complex. Cohesin crossing category-specific binding factors and cohesin-cohesin crossing rates were set at 0.45 and 0.15 crossings per second, respectively, because cohesin occupancy at CTCF binding sites is typically more pronounced. The ensemble of chromatin conformations resulting from these simulations were used to generate pair-wise interaction matrices among the beads of each polymer, allowing for direct quantitative comparisons between the model and experimental Micro-C data.

## Acknowledgements

We thank the Söding lab (MPI-NAT) for their feedback, Alexis Bel for his help in data processing, and all the Papantonis lab members for discussions. This study was supported by the German Research Foundation (DFG) via the Priority Program 2202 (project 422389065 awarded to A.P.), by the Lower Saxony Ministry for Research and Culture (MWK) via the SPRUNG (project 76211-1267/2023 awarded to A.P.) and BEREIT programs (project 2019-00298 awarded to A.P.). Y.Z. was supported by the IMPRS Molecular Biology program. M.B. is supported by the Priority Program 2202 ‘Accelerator Award’ and the Else-Kroener-Fresenius Medical Scientist program of the UMG.

## Author contributions

Y.Z. and M.B. developed code and performed computational analyses; A.P. conceived the project and secured funding; Y.Z., M.B. and A.P. co-wrote the manuscript.

## Declaration of interests

The authors have no competing interests to declare.

## Supplementary Figures

**Figure S1.**
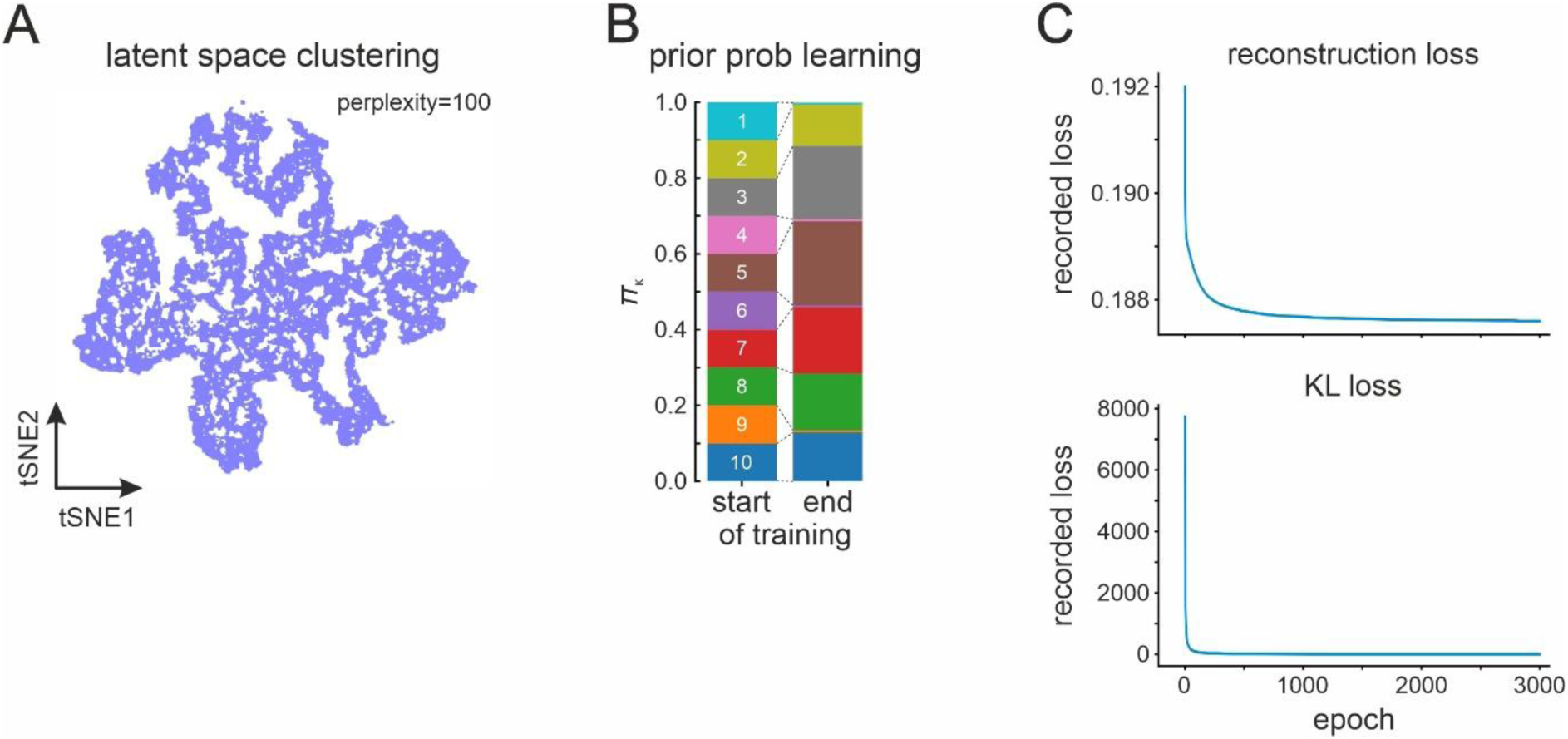
LoopBin training features. A, GMM collapses to the unit gaussian in the latent space when μ*k* and θ*k* learning is omitted from the pre-training phase (t-SNE of latent space; perplexity=100). B, LoopBin conststently empties redundant loop classes. C, Loss function optimization during 3000 epochs of the training phase.

**Figure S2.**
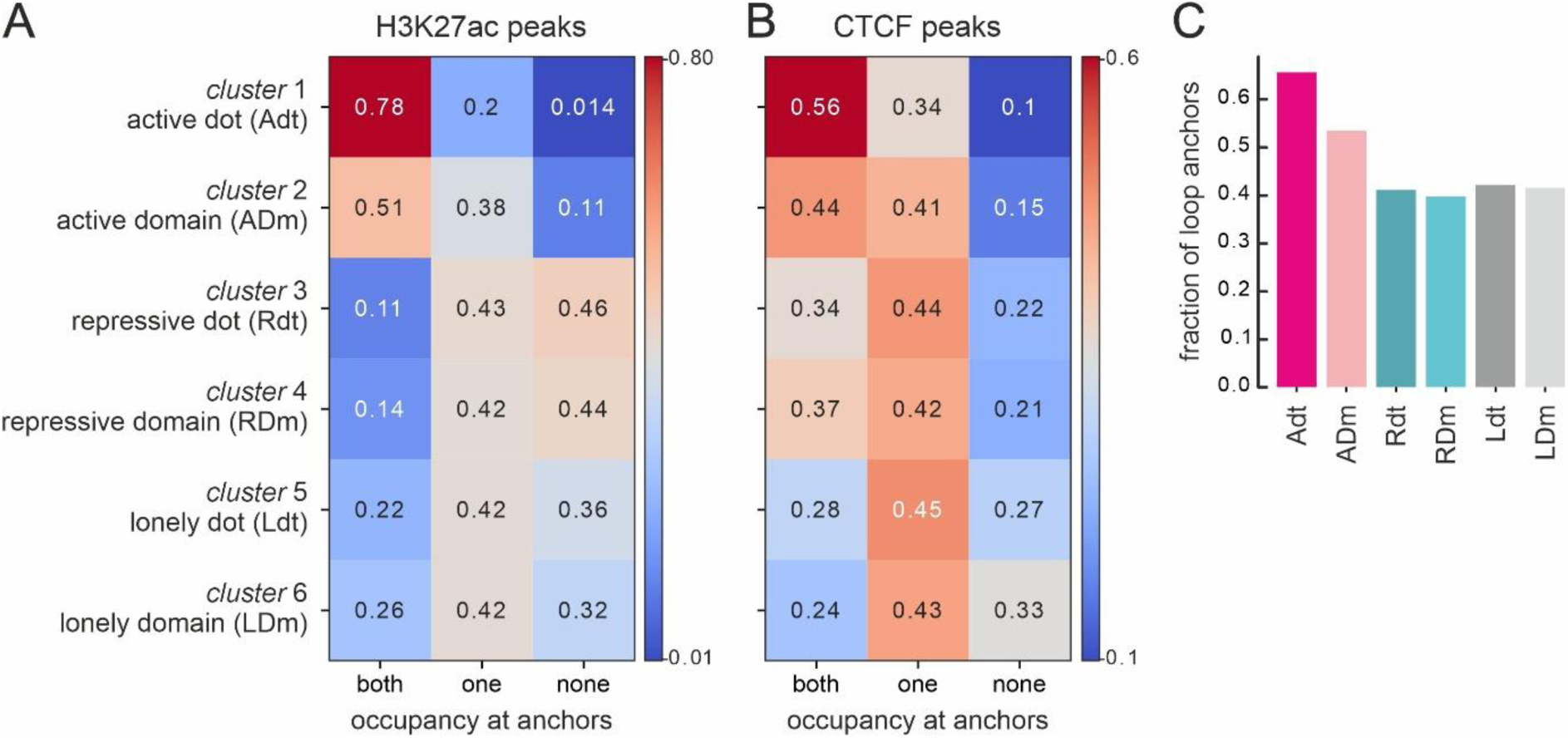
LoopBin classification compared to a manual annotation. A, LoopBin loop classes crossed with a manual annotation based on the presence of H3K27ac peaks at both, only one or at none of the loop anchors. B, As in panel A, but for CTCF peaks at loop anchors. C, Number of promoters positioned in the 10 kbp around loop anchors of each Loopbin class.

**Figure S3.**
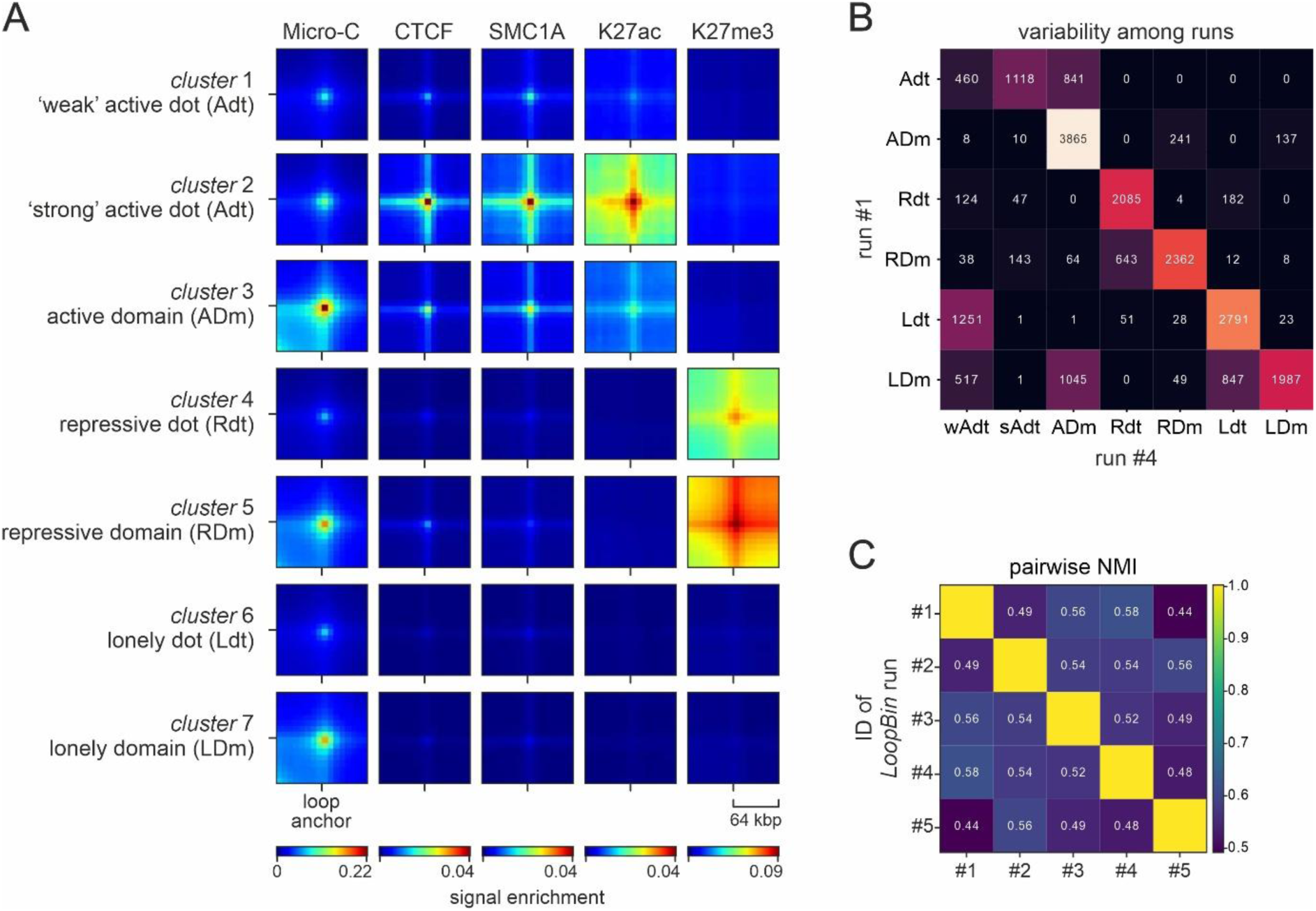
LoopBin classification reproducibility. A, Due to the continuity of input data, LoopBin may split one class into subcategories. In this example, split the Adt class into weak and strong subclasses based on the overall strength of H3K27ac signal. All other identified classes remain essentially unchanged. B, Heatmap showing the number of loops shared between the LoopBin classes from panel A and the six classes shown in Fig 1. C, Heatmap showing normalized mutual information (NMI) values for five independent LoopBin runs.

**Figure S4.**
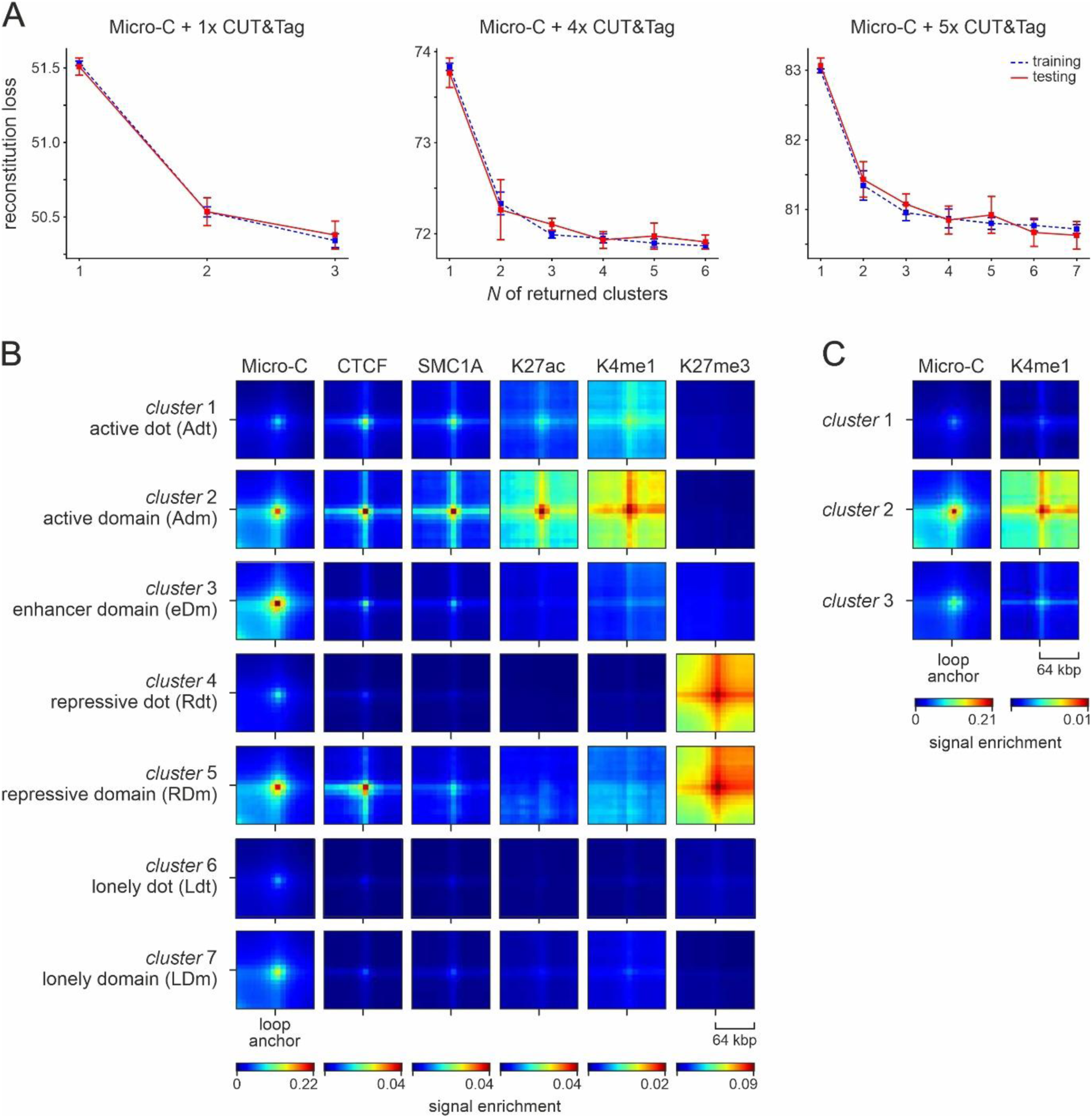
LoopBin classification is influenced by the number of epigenomic tracks available. A, Reconstruction and regularization error in LoopBin when provided with a different number of epigenomic (CUT&Tag) datasets and returning an increasing number of classes. B, LoopBin classification upon addition of H3K4me1 data to the input produces an additional class to those identified in Fig 1. All other classes remain essentially the same. C, As in panel B, but using only Micro-C and H3K4me1 data as input.

## Notes

### Competing Interest Statement

The authors have declared no competing interest.

